# Investigation of CMA Substrate Interactions with Hsc70 Reveals Potential Auxiliary Binding Sites

**DOI:** 10.1101/2025.01.30.635767

**Authors:** Devid Sahu, Nidhi Malhotra

## Abstract

Chaperone-mediated autophagy (CMA) is a selective lysosomal degradation pathway crucial for cellular homeostasis. Its dysfunction is linked to cancer and neurodegenerative disorders. Hsc70 mediates CMA selectivity by recognizing proteins with ‘KFERQ-type’ sequences, but the structural basis for substrate recognition and the substrate-binding mechanism of Hsc70 remain unclear. This study integrates literature, bioinformatics, AlphaFold 3 predictions, and 44 μs of molecular dynamics (MD) simulations to elucidate the lid domain interactions of Hsc70 with fifteen experimentally verified CMA substrate segments. Structural and thermodynamic residue-level analyses suggest stable binding of substrates in two distinct sites. The first site resides in a hydrophobic cavity formed by alpha helix α1 and α3 in the lid domain of Hsc70, with complex stabilization primarily driven by hydrophobic residues within ‘KFERQ-type’ motif and its flanking regions. The second stable site was observed to be canonical hydrophobic cleft in Hsc70, with complex stabilization primarily driven by hydrophobic residues within ‘KFERQ-type’ motif as well as the through water-mediated interactions by the lid domain. The robustness of our findings is confirmed by consistent convergence across force fields, substrates, and supporting literature, all identifying the same site. We propose these sites may facilitate upstream or downstream recognition and activation events.

**Author Summary:** Cells must constantly remove damaged or unnecessary proteins in order to stay healthy. One important pathway that performs this task is chaperone-mediated autophagy. When this pathway fails, it has been linked to conditions such as cancer and neurodegenerative diseases. A key player in this process is the protein Hsc70, which recognizes short sequence signatures in target proteins and directs them for degradation. However, how Hsc70 physically recognizes these signatures has remained poorly understood. In this study, we investigated how Hsc70 interacts with a range of confirmed target protein segments. We combined information from prior experimental studies, with computational structure prediction and extensive molecular simulations to explore these interactions in detail. Our results indicate that Hsc70 can bind target proteins at two distinct regions, each stabilized mainly by hydrophobic amino acids within and/or around the recognition signal, with additional contribution from water-mediated interactions in the latter site. Notably, these binding modes were consistent across different targets and computational conditions. Together, our findings provide a clearer structural picture of how selectivity is achieved in this degradation pathway. This improved understanding may help guide future studies on how chaperone-mediated autophagy is regulated and how it becomes disrupted in disease.

## Introduction

Chaperone-Mediated Autophagy (CMA) is a selective degradation process with diverse roles in maintaining cellular homeostasis, regulating immune response, influencing aging, and contributing to the pathogenesis of various diseases, including cancer[1–5]. The unique feature of selectivity distinguishes it from other types of autophagy. The selectivity is rendered by a heat shock cognate protein, Hsc70, that recognizes only a cohort of selective proteins called substrates and only such proteins are amenable to degradation by CMA. Despite its well-established function in recognizing and binding substrate proteins, the precise molecular details governing the interaction between Hsc70 and its substrate proteins remain largely unknown. Sequence-based designation of the substrate requires presence of ‘KFERQ-like’ (CMA-targeting) motifs in them[6–9]. This motif typically includes one negatively charged residue (D or E), one or two positively charged residues (K or R) and one or two hydrophobic residues (I, L, V, or F) in any order along with a glutamine (Q) at either end of the motif. The “KFERQ-like” motif can be turned on or off by chemical changes to proteins i.e post-translational modifications (ubiquitination, phosphorylation or acetylation). KFERQ-like motifs without any modifications are called canonical motifs[10–13]. Currently, ∼98 protein substrates are known for CMA, of which 76 substrates have been identified in humans[13–20]. The challenge lies in deciphering how Hsc70 achieves selectivity for substrate proteins in a cellular environment where multiple proteins bear similar recognition motifs. Particularly, what are the structural fingerprints that label a protein as a CMA substrate, how does Hsc70 recognize those fingerprints, what is the substrate binding site in Hsc70? This gap in understanding has significant implications, as substrate recognition is the cornerstone of CMA’s selectivity, influencing various cellular processes and disease outcomes.

Hsc70 is a constitutive member of the Hsp70 family that shares a conserved structural architecture comprising of a nucleotide binding domain (NBD), and a substrate binding domain (SBD). The SBD is sub-divided into SBDβ, SBDα (lid) and a C-terminal extension[21,22]. Most of the existing literature exclusively utilized short peptides as model substrates to develop the current understanding of chaperone-substrate binding[23–26]. Crystal structure of a model peptide NRLLLTG (NR) in complex with DnaK suggested that the peptide binds in an extended conformation in a canonical hydrophobic cleft within SBDβ[26]. Peptides constituting CMA-targeting motifs of substrate protein ‘Tau’ were experimentally shown to bind independently and did not compete with model peptides for binding at SBDβ[27]. In the same study, NMR experiments have shown that these KFERQ-like peptide motifs were unable to bind Hsc70’s SBD that contained only SBDβ. This suggests that either the substrate protein containing KFERQ-like motifs bind at a site other than the canonical hydrophobic binding groove or the binding of KFERQ-like motifs is weaker than the model peptide at the same site with possibilities of existence of secondary binding sites. It also points to the role of SBDα in binding. Additionally, SBDα is the sequentially divergent region amongst Hsp70 orthologs and residues being involved in binding at this site can account for the differential binding of tau with Hsc70 and Hsp72[28].

There are several bottlenecks towards gaining a comprehensive understanding of the interactions between Hsc70-substrate complex. Firstly, the atomistic structure of Hsc70 is unknown and the exact binding site is unidentified. Additionally, consistent with the principle applicable to chaperones in general, its binding with the client protein must be strong enough to prevent the spontaneous detachment of the bound protein yet not so strong to enable its detachment for subsequent downstream events. Thus, these chaperone-client complexes are considered as “fuzzy” complexes. The binding interfaces constitute formation of transient, dynamic, and nonspecific interactions, capable of modulating the conformational equilibrium, which are difficult to capture experimentally. The groundbreaking deep learning approach of AlphaFold 3 predicts structures of proteins and their complexes by leveraging evolutionary and structural data[29]. Combined with MD simulations, which model atomic interactions and dynamic motions, it provides a powerful framework to analyze these transient and dynamic interactions, overcoming experimental limitations. In the current study, we have utilized an integrated approach combining information from existing literature, molecular modelling, AlphaFold 3 and molecular dynamics simulations to predict Hsc70-substrate binding interactions. Our findings suggest that the residues adjacent to the CMA-targeting motifs in the same secondary structure element stabilize the binding interactions between SBDα of Hsc70 and the substrate. The analysis on these extended CMA-targeting motifs in fifteen human substrate proteins reveal common binding site on SBDα of Hsc70 encompassing residues Tyr545, Asn548, Thr552, Leu558, Lys561, Glu600, Ile607, Leu610, and Tyr611. The site consists of hydrophobic interior with hydrophilic residues flanking the central pocket. The use of various force fields, different substrates, and supporting evidence from existing literature-all converging on the same site, validates the robustness and success of our study. The functional implications of the binding may not be to render selectivity to the CMA pathway since the KFERQ pentapeptide is necessary and sufficient for binding[30], but they might contribute to facilitating upstream or downstream recognition or activation events. The study opens the door to further exploration in this direction. Additionally, we observed stable and selective binding of CMA-targeting motifs to the canonical site in SBDβ, with major contributions from the hydrophobic amino acids of the motif, along with significant interactions involving residues in SBDα of Hsc70.

## Results and Discussion

### Adjacent Residues contribute to the Stability of ‘KFERQ-like’ Motif Binding to Hsc70’s lid domain

The binding of Hsc70 to CMA substrates has primarily been studied at the sequence level to date, with the well-known recognition of ‘KFERQ-like’ motifs in substrates involved in binding. At the structural level, evidence from multiple sources in literature suggests potential role of residues at the lid domain/SBDα of Hsc70[27,28,31–33]. To understand the structural basis for substrate recognition and the substrate-binding mechanism to SBDα of Hsc70 (if any), we compiled a list of CMA substrates and analyzed their interaction with SBDα of Hsc70, focusing on the recognition of ‘KFERQ-like’ motif.

A comprehensive list of experimentally validated CMA substrates in humans was collated. Using the KFERQ Finder bioinformatics tool, both canonical KFERQ motifs and KFERQ-like motifs generated by post-translational modifications (PTMs) were identified[13]. Fifteen substrates containing a single canonical motif were selected for analyzing the recognition and binding site in Hsc70 SBDα. The selected substrates are: RND3(Rho-related GTP-binding protein RhoE), Abeta(Amyloid-beta precursor protein), Annexin A1, Annexin A2, Annexin A6, GBA1(Lysosomal acid glucosylceramidase), Cyclin D1(G1/S-specific cyclin-D1), PEA15(Astrocytic phosphoprotein PEA-15), HBB(Hemoglobin subunit beta), CHK1(Serine/threonine-protein kinase Chk1), α-Synuclein, Annexin A4, Prion protein(Major prion protein), PKM(Pyruvate kinase PKM), and PARK7(Parkinson disease protein 7). The list along with their canonical motifs are enlisted in S1 Table. AlphaFold 3 was utilized to predict the structure(s) of the complex(es) between substrate’s ‘KFERQ-like’ motif and SBDα of human Hsc70. Fig 1A depicts the residue-level interaction profiles of Hsc70 with different CMA substrate segments. Majorly, three distinct sites were observed (site-1 is colored light steel blue, site-2 as pink and site-3 as peru), Fig 1B. Few representative protein complexes from each site (RND3, Abeta and Annexin A1 from site-1, CHK1, and PKM from site-2 and PARK7 from site-3) were subjected to all-atom MD simulations. The stability of the binding site was checked by plotting the minimum distance between the substrate and the receptor, Fig 1C and S1A Fig. In all cases, the predicted binding site demonstrated structural instability during the simulations, failing to maintain its integrity throughout the trajectory. Consequently, the substrate exhibited a loss of association with the putative binding pocket and migrated away from the predicted site. This dislocation suggests a potential need to refine the molecular environment to accurately replicate the native binding dynamics.

**Fig 1.**
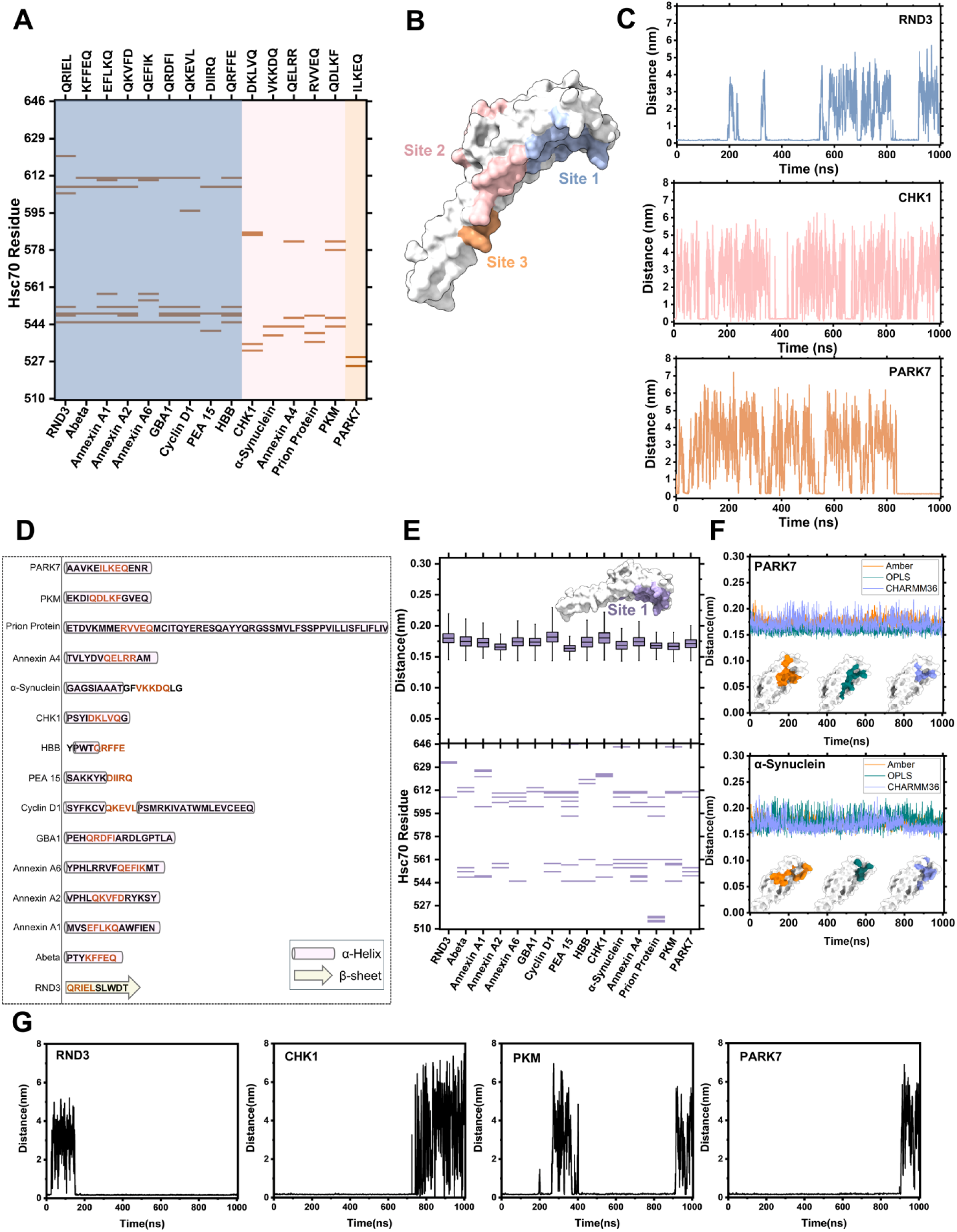
Residues adjacent to the CMA-targeting motif stabilize its binding with the lid domain of truncated Hsc70. (A) Heatmap representing the residues of Hsc70 lid domain interacting with ‘KFERQ-like’ motifs in AlphaFold 3 predicted complexes for 15 CMA substrates. Three different sites were observed (site-1 in light steel blue, site-2 in pink, and site-3 in peru). (B) 3D surface representation of the lid domain of Hsc70 depicting the three predicted binding sites. The loop region in the lid domain is not shown for clarity. (C) Minimum distance between Hsc70 lid domain and CMA substrate peptides with only ‘KFERQ-like’ motif. It represents structural instability of the complex during the course of MD simulations. (D) Schematic representation of secondary structure of CMA-targeting motifs along with adjacent amino acid residue of 15 CMA substrates that were taken for binding site predictions (termed as ‘extended CMA-targeting motifs’). (E) The minimum distance plot between the Hsc70 lid domain and the extended CMA-targeting motifs (Top panel), along with heatmap representing the interacting residues of the two (Bottom panel), averaged over last 200 ns. Surface representation of Hsc70 lid domain with the interacting side highlighted in purple (loop region is not shown for clarity). The binding site remained stable throughout the simulations. (F) Minimum distance plots of Hsc70 lid domain and extended CMA-targeting motifs in PARK7 and α-Synuclein using AMBER, OPLS, and CHARMM 36 force fields. A value below 2.5 Å indicates stability in each force field. (G) Minimum distance between Hsc70 and RND3, CHK1, PKM and PARK7 peptide in the complexes. The complexes were generated by removing the adjacent residues in CMA substrates after they had participated in forming a complex with the motif and the chaperone. None of the complexes remained stable.

Since the ‘KFERQ-like’ motif alone does not maintain sufficient stability in complex with Hsc70, we investigated the potential contribution of adjacent residues in stabilizing the motif’s interaction with the chaperone, as their presence may be essential for effective binding. Extensive stabilization by additional residues outside the consensus sequence motifs have also been previously observed for LIR motifs[34,35]. LIR motif is flanked by C-helix residues that cause additional stabilization. To define ‘adjacent’ residues in our study, the subset of shortlisted substrate proteins is analyzed for the existence of secondary structural elements in ‘KFERQ-like’ sequence motifs (Fig 1D). It was observed that the motif forms a part of alpha-helical conformation in ten proteins. In only one and four proteins, on the other hand, the motif exists in beta-sheet and loop respectively. Those existing in the loop have flanking residues in alpha-helical conformation. Adjacent residues are defined as variable-length stretches of residues flanking the ‘KFERQ-like’ motif on either side that adopts same (in case of alpha-helix and beta-sheet) or adjacent (in case of loop) secondary structure component as residues of the motif. These residues along with the motif, hereafter referred as ‘extended motifs’ were used for exploring the binding site between the substrate proteins and Hsc70 SBDα.

AlphaFold 3 was utilized to predict the structure of the complex between the peptides with extended motifs and the lid domain of Hsc70. To check the stability of these complexes, each of them were subjected to 1 *µ*s of all-atom MD simulations, generating a total of 15 *µ*s of data. Interestingly, compared to the previous case wherein only ‘KFERQ-like’ motifs were explored, the extended motifs demonstrated structural stability of the complex. The minimum distance between the substrate and the receptor protein remained below 2.5Å, indicating stable binding during the timescale of MD simulations (Fig 1E top panel and S1B Fig.). The residues of Hsc70 involved in the interaction with the residues of the substrate during the last 200 ns of the simulations are illustrated in Fig 1E (bottom panel). It was observed that the simulations stabilize all the proteins to site 1 of Hsc70. To investigate whether the observed binding stability was an artifact of a particular force field or a general phenomenon, we performed simulations of the two of the complexes generated by AlphaFold 3 (PARK7 and α-Synuclein) using multiple force fields (CHARMM36, AMBER, and OPLS) with varying parameterization schemes. It was observed that the substrates are bound throughout the trajectory (Fig 1F). Overall, across all tested force fields, the binding remained intact, suggesting that the interaction is robust and not specific to any single force field. This consistency enhances confidence in the accuracy of the binding site prediction and its relevance under physiological conditions. Furthermore, the site is in agreement with the previous proposed model based on cross-linking experiments between human Hsc70, yeast ssa1p and α-Synuclein[33].

To evaluate the contribution of adjacent residues to the stability of the ‘KFERQ-like’ motifs in the bound state, we removed the adjacent residues after they had participated in forming a complex with the motif and the chaperone i.e. adjacent residues flanking the motif were removed from the bound complexes in four of the previous runs (RND3, CHK1, PKM and PARK7) and the resulting complexes were subjected to MD simulations. The time evolution of the distance between substrate and receptor was monitored thereafter to check the stability of the complexes, Fig 1G. None of the complexes remained stable at the same site. The substrate disengaged from the proposed binding region and relocated to alternate positions. These findings indicate that the binding interactions may not have been sufficiently robust. Taken together, our data suggests that additional residues adjacent to ‘KFERQ-like’ motif appear to play a crucial role in stabilizing the complex with the lid domain of Hsc70 under the conditions studied.

Since it is well-established that the pentapeptide motif in substrates is both necessary and sufficient for performing its CMA activity[30], we believe that the binding of CMA substrates in this region may not have direct implications in rendering selectivity to the process. Instead, the observed interactions point toward the presence of an additional binding site, which may play a role in upstream events such as substrate recruitment or in downstream processes like substrate transfer to LAMP2A. Whether and how the site could impact substrate triage, retention, or handoff to LAMP2A requires further experimental investigation. Nonetheless, the identified site remains robust, with supporting experimental evidence from the literature[32,33].

### Extended CMA-targeting motifs in substrate proteins have selective affinity to hydrophobic cavity formed by helix α1 and helix α3 in SBDα of Hsc70

To delve deeper into the binding site and interaction pattern, the binding pocket in Hsc70 and the nature of interacting residues were identified. Fig 2A depicts a chord plot representing the residues involved in interaction. Following an extensive analysis of the trajectories, we identified a set of receptor residues that consistently participated in interactions across majority of the simulated complexes. These residues include Tyr545, Asn548, Met549, Asp555, Lys557, Leu558, Lys561, Glu600, Asn604, Ile607, Leu610, Tyr611, which were involved in stabilizing the binding through van der Waals interactions. Notably, Ile607 and Tyr611 emerged as critical hotspots, forming persistent interactions with the peptide in over 8 of the complexes. The conservation of these interaction patterns across the majority of systems underscores their potential significance in maintaining the binding and highlights their role in the structural integrity of the receptor-substrate complex. Overall, it was observed that interacting residues belong to helix *α1* and helix *α3* of SBD*α* in Hsc70 (Fig 2B).

**Fig 2.**
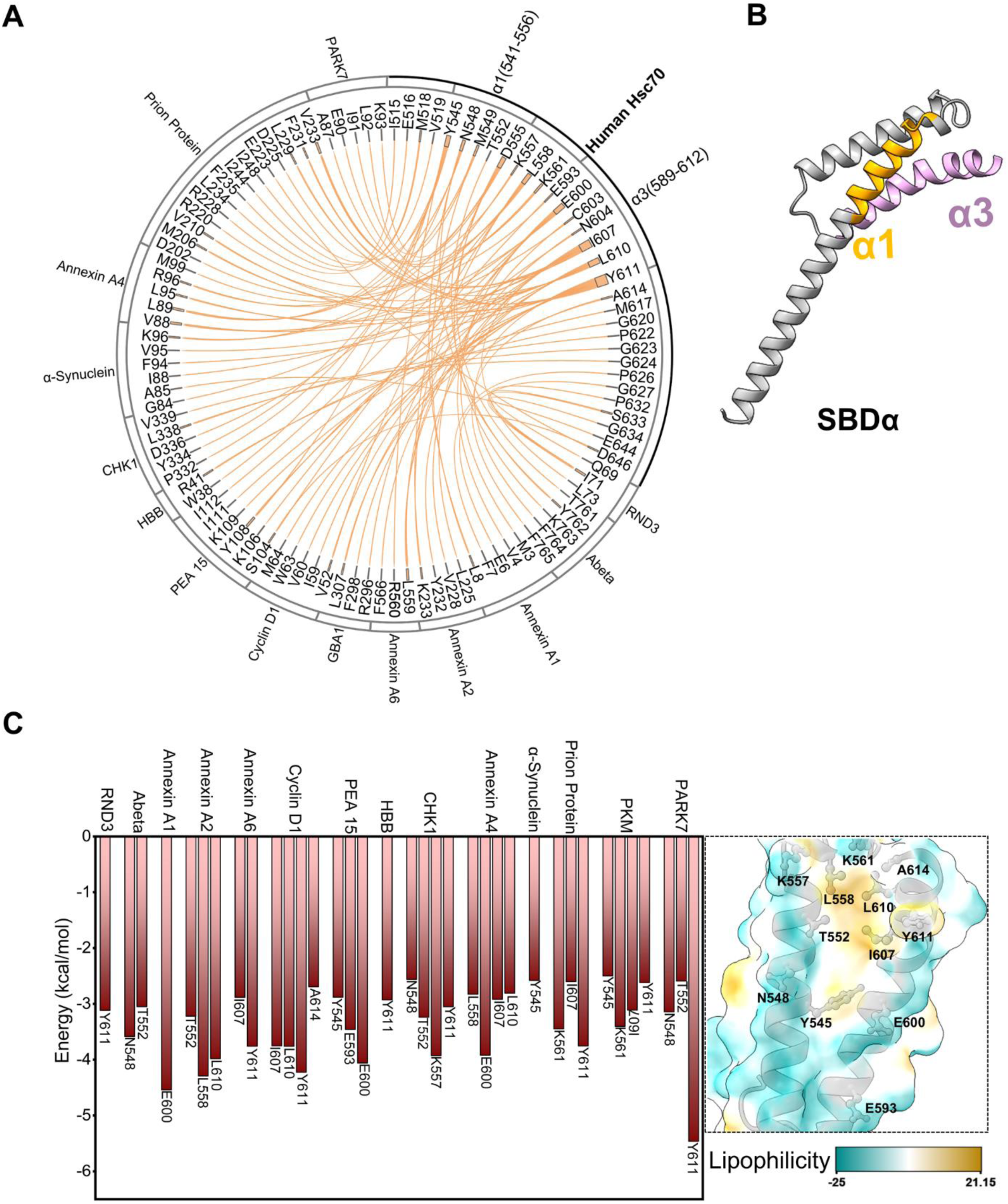
Extended CMA-targeting motifs selectively bind to the hydrophobic cavity formed by helix α1 and helix α3 in the lid domain of truncated Hsc70. (A) A chord diagram representing the residues of Hsc70 lid domain interacting with extended CMA-targeting motifs from 15 CMA substrates, where the thickness of the node for each respective residue represents the frequency of interactions. Residues of helices α1 and α3 are extensively participating in interactions with the substrate segment. (B) 3D cartoon representation of the lid domain of Hsc70 depicting the helices α1 and α3 which are involved interactions with the substrate segment. The loop region in the lid domain is not shown for clarity. (C) Residue-wise decomposition of binding energies of Hsc70 residues with the CMA substrate peptides is shown in the left panel. The bar plot of binding energies of residues with values less than −2.5 kcal/mol (calculated using MMGBSA) are shown for clarity. Right panel showcases lipophilic representation of the binding pocket in Hsc70. Key interacting residues are depicted in ball-and-stick representation.

The contribution of each Hsc70 residue towards stabilizing the binding interactions were analyzed by performing free energy calculations using MM-GBSA (Molecular Mechanics-General Born Surface Area) approach. Fig 2C showcases the binding energies of these residues with values more negative than −2.5 kcal/mol. Most of the residues are consistent with our occupancy analysis above. The common residues of occupancy and binding energy analysis include Tyr545, Asn548, Thr552, Leu558, Lys561, Glu600, Ile607, Leu610, and Tyr611. We propose the significance of these residues in rendering stabilization to the complex. Further, the identified residues in Hsc70 encompassing the site contribute to the formation of a cavity with hydrophobic interior and hydrophilic exterior. The site can accommodate the side chains of hydrophobic amino acids of the peptide segment.

### Extended CMA-targeting motifs in substrate proteins display promiscuous orientation in the binding hydrophobic cavity of SBDα

The stabilizing interactions between the substrate and chaperone within the selective binding site were examined in greater detail. Interestingly, despite the substrate consistently associated with the designated binding pocket, no common binding orientation of the substrate was identified. Our simulations suggested that the substrate(s) adopt different binding modes in each simulation, leading to variable contact patterns with the binding site, S2A Fig. A comprehensive analysis of interaction patterns revealed a lack of reproducible stabilizing contacts across simulations of different substrates. Quantitative analysis, including calculating the contact frequencies and interaction energy underscores this observation (Fig 2A and S2B Fig. respectively). Nevertheless, in all the simulations, hydrophobic amino acids were observed to have more negative interaction energy. The minimum distance between key interacting pairs of Hsc70 and the substrate(s) is depicted in S3 Fig. These interactions were stable with the value ranging within 4Å. However, no consistent pair was identified in all the simulations. This variability indicates that, while the substrate binds selectively to the hydrophobic pocket, the mode of binding demonstrates considerable flexibility. The absence of a consistent interaction pattern suggests a promiscuous nature of substrate-chaperone binding to this site.

When simulations were performed using different force fields, this dual characteristic of binding was consistently observed: while the substrate exhibited high specificity for the binding site across all force fields, the nature and dynamics of individual interactions varied significantly (S4 Fig.). Certain force fields emphasized stronger hydrophobic contacts, while others resulted in a more diverse range of transient interactions. These differences highlight the role of force field parameters in capturing the flexibility of substrate binding modes, suggesting that the observed promiscuity may be an inherent property of the system rather than an artifact of a specific simulation setup. Overall, the data suggest that substrate binding to the observed site in SBDα is promiscuous and does not by itself account for substrate selectivity. Instead, our data reveals a potential secondary site in the lid domain. The functional implication of this site requires further experimental investigation. The observed site is however consistent with previous cross-linking experiments[32,33].

### Stable recognition of CMA-targeting motifs requires the closed conformation of full-length Hsc70

Owing to the obtained promiscuous nature of binding of CMA-targeting motifs with SBDα of Hsc70, we next investigated the binding of the motif with full-length Hsc70 (along with ADP) to reason its selective behavior in CMA. AlphaFold 3 predicted the binding site to be the canonical hydrophobic cleft in SBDβ for all the 15 tested substrates. Interestingly, 4 of them were predicted to be bound to the ADP-bound closed form of Hsc70 while remaining 11 were bound to its open form (Fig 3A-B and S5 Fig.). All the closed-form complexes (Abeta, Annexin A1, PEA 15 and HBB), along with six representative open-form (RND3, α-Synuclein, CHK1, PKM, Annexin A6 and PARK7) complexes were subjected to MD simulations of 1 μs each to check their stability. The analysis of the time evolution of the minimum distance between the substrate and Hsc70 suggested that complexes bound to the closed form were stable throughout the time scale of our simulations, while the open forms exhibited pronounced fluctuations (Fig 3C and D). This suggests that efficient recognition of KFERQ-like motifs requires conformational coupling between the nucleotide-binding domain (NBD) and SBDβ, and/or the presence of additional structural elements such as the interdomain linker or SBDα/lid. Consistent with our observations, the importance of ATP allostery of Hsc70 in regulating the interaction has also been shown experimentally[36].

**Fig 3.**
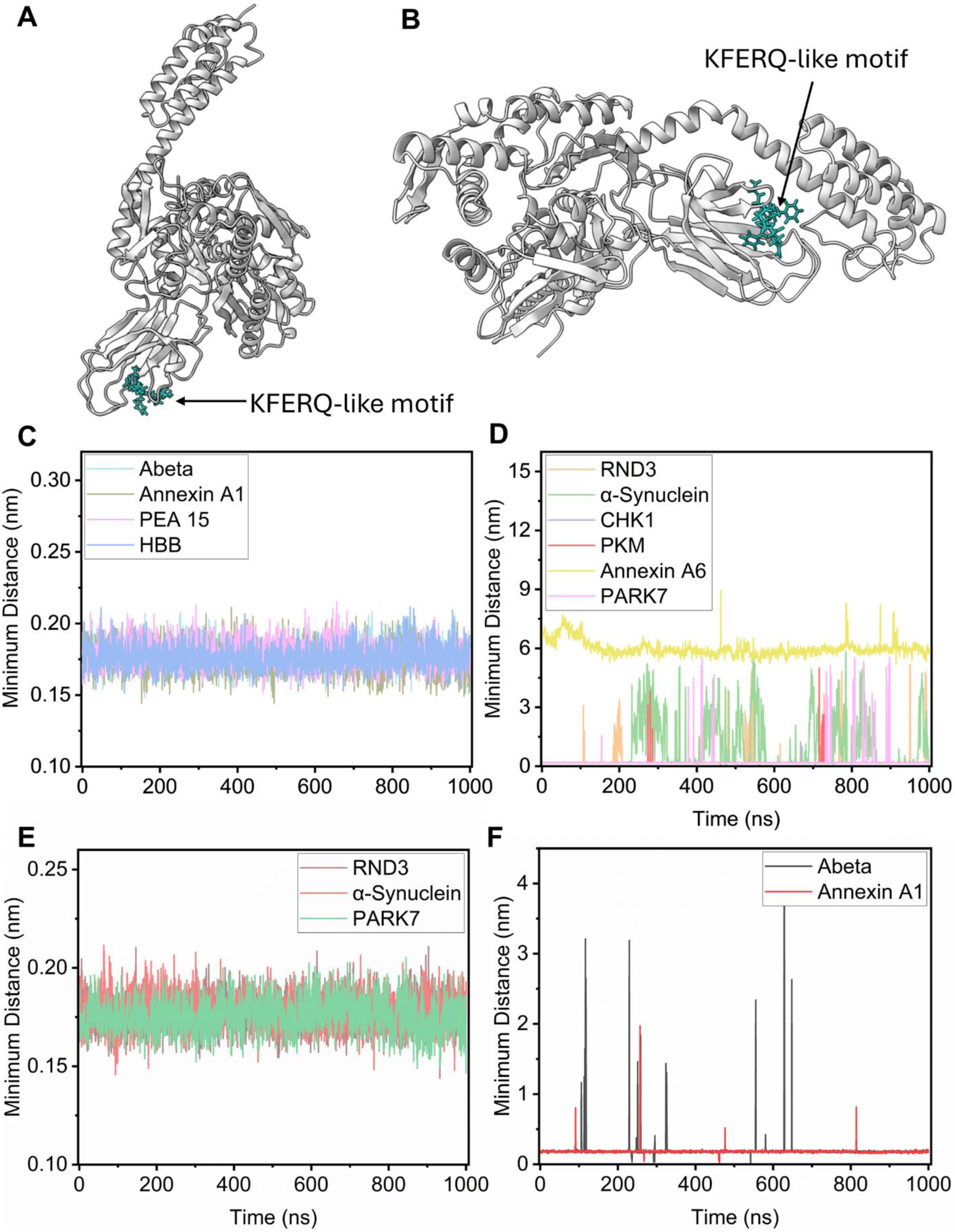
CMA-targeting motifs bind to full-length Hsc70 in its closed conformation, where the lid domain plays an essential role in stabilizing the interaction. Representative snapshot of AlphaFold 3 (AF3) predicted conformation of Hsc70 (dark gray)-substrate (light sea green) complex with Hsc70 in (A) open conformation and (B) closed conformation. Time evolution of the minimum distance between CMA-targeting motif and Hsc70 in AF3 predicted substrate-Hsc70 complexes wherein Hsc70 is in (C) closed formation, (D) open conformation, (E) open conformation initially but were converted to the closed conformation keeping initial contacts intact, F) closed conformation initially but the lid domain (residues 510–646) was removed before subjecting it to MD simulations.

The unstable open form of Hsc70-substrate complexes in our MD simulations raises the question of whether the instability arose from weak substrate-chaperone interactions themselves or from the absence of the lid domain. To disentangle these possibilities, we preserved the substrate-chaperone contacts from the three of the representative AF3-predicted open forms (RND3, α-synuclein, PARK7) and converted the complexes into closed conformations. Each system was then subjected to 1 μs of MD simulations. In all three cases, the substrate remained stably bound throughout the trajectories (Fig 3E), supporting the view that lid closure is critical for stable substrate engagement.

Having established that lid closure promotes stable substrate engagement, we next examined the effect of lid/SBDα removal. To this end, we truncated the SBDα from two representative closed-form complexes (Abeta and Annexin A1), thereby mimicking isolated SBDβ bound to the KFERQ-like motif while retaining the previously established interactions. Both systems were subjected to 1 μs of all-atom MD simulations. In both cases, the CMA-targeting peptide failed to remain in a stable pose over the course of the simulation (Fig 3F). These results, together with our earlier observation that full-length Hsc70 stably accommodates the motif, are consistent with experimental reports that KFERQ motifs fail to bind isolated SBDβ constructs[27]. Collectively, these results highlight the importance of investigating substrate recognition in the context of the full-length chaperone.

### CMA-targeting motifs selectively bind the canonical hydrophobic cavity of SBDβ, preserving backbone alignment despite side-chain registry shifts

To gain mechanistic insights into how Hsc70 closed conformation stabilizes substrate binding, we analyzed the residues and interaction sites in full-length Hsc70 involved in recognizing CMA-targeting motifs (Fig 4A). Ala 406, and Thr 429 in Hsc70 were observed to have occupancies exceeding 90% of the simulation time, across all the tested substrates. All of the stable CMA substrate complexes in our study were bound to the canonical hydrophobic cavity within SBDβ (residues 394-509 of Hsc70), Fig 4B. NRLLLTG peptide has also been previously reported to be nestled in the same groove of DnaK[37] and Hsp70[38]. The binding site in these proteins is organized into five distinct pockets (−2,-1,0,1,2) that accommodate the extended peptide backbone, while the helical lid of the SBD folds over to enclose the substrate. The central pocket designated as position “0”, is highly specific and primarily accommodates hydrophobic residues. The remaining pockets are numbered relative to the central pocket. For the NRLLLTG peptide, it is reported that the central leucine residue fits optimally into the pocket 0. The flanking pockets are less specific and can accommodate diverse substrate side chains, thereby conferring binding promiscuity.

**Fig 4.**
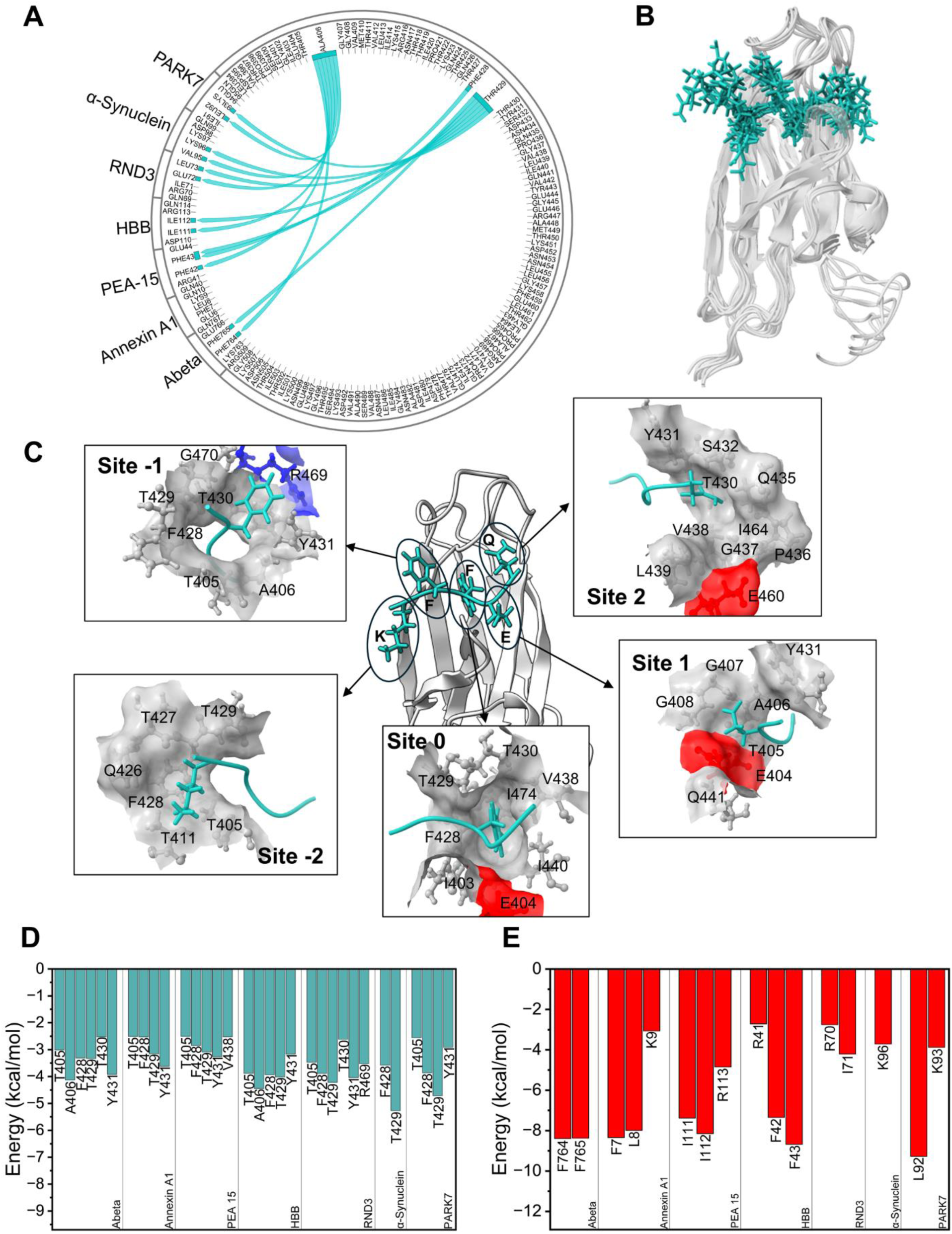
CMA-targeting motifs bind to the hydrophobic cleft of the SBDβ domain in the closed conformation of Hsc70, maintaining the backbone registry and being stabilized primarily by hydrophobic residues. (A) Chord plot representing the interacting residues of Hsc70 SBDβ domain and the CMA-targeting motif of seven CMA substrates in which Hsc70-substrate complex is stable during 1μs of MD simulations. The thickness of the node for each respective residue represents the frequency of interactions For Clarity, residues with interaction occupancy more than 90% are shown. (B) Overlay of the CMA-targeting motifs (light sea green) in these seven CMA substrates, when bound to closed conformation of full-length Hsc70 (dark gray). For clarity, only SBDβ of Hsc70 is shown. (C) Snapshot of CMA-targeting motif of Abeta (light sea green in stick representation) bound to Hsc70 (dark grey in cartoon representation) complex at 1μs of MD simulation data. Again, for clarity, only SBDβ of Hsc70 is shown. Surface representation of five distinct pockets within hydrophobic cleft of SBDβ in Hsc70 accommodating five residues of the motif are shown alongside. Pocket residues are explicitly labeled and colored gray, blue, or red to represent hydrophobic, basic, and acidic amino acids, respectively. (D) Residue-wise decomposition of binding energies of Hsc70 residues with the CMA substrate peptides. (E) Residue-wise decomposition of binding energies CMA substrate peptides with Hsc70. For clarity, only the bar plots of binding energies of residues with values less than −2.5 kcal/mol (calculated using MMGBSA) are shown in panels (D) and (E).

In line with these observations, we analyzed the placement of the pentapeptide residues within the five pockets. A representative example of a KFERQ-like motif from Abeta interacting with pocket residues is shown in Fig 4C. Our results suggest that the central binding site (site 0) in Hsc70 comprising residues Ile 403, Glu 404, Phe 428, Thr 429, Thr 430, Val 438, Ile 440, and Ile 474 serves as the most selective site stabilizing hydrophobic residue(s) of CMA-targeting motifs in all seven stable complexes in MD simulations (Closed form of Hsc70 with CMA-targeting motifs of (i) Abeta, (ii) Annexin A1, (iii) PEA-15, (iv) HBB, along with AF3-predicted open form Hsc70 complexes that were converted into complex with Hsc70 in closed conformations: (v) RND3, (vi) α-Synuclein, and (vii) PARK7). The remaining pockets, by contrast, are less specific, accommodating both hydrophobic and polar residues of CMA-targeting motifs and contributing only minor interactions. Binding free energy analysis using MM-GBSA further supports significant contribution of residues involved in site 0, as depicted in Fig 4D. Collectively, our findings that nonpolar interactions provide the major energetic contribution to binding are consistent with prior reports for DnaK[39].

The canonical binding groove of Hsp70 is well known to exhibit registry plasticity and a lack of strict directional bias, as both forward and reverse orientations of peptide binding have been reported previously[40]. Additionally, comparative structural studies have revealed a one-residue registry shift between bacterial DnaK and human Hsp70/BiP bound to similar NR peptides, indicating that even canonical motifs can adopt alternate alignments. In this context, our structural modeling of the KFERQ-like motifs revealed a registry nearly identical to that observed for Hsp70-bound NRLLLTG, with the hydrophobic residue fitting into the 0 subsite, with a specific preference to leucine residue. Other hydrophobic residues like isoleucine, and phenylalanine were also observed to bind at site 0 (S2 Table). Energy analysis suggests that these hydrophobic residues are the major contributors of the interaction energy (Fig 4E).

Interestingly, even though side-chain positioning within the substrate-binding groove may vary i.e., different residues occupy the hydrophobic pockets at subsites 0, ±1, and ±2, but the backbone registry remains conserved, (Fig 4B, S6 Fig.). Irrespective of the substrate, backbone residues Ala406 and Thr429 were observed to form hydrogen bonds with the backbone of the CMA-targeting motif. This structural feature was robust across independent simulations and aligns with prior structural observations in both bacterial DnaK and mammalian Hsp70, where registry shifts at the side-chain level did not disrupt the backbone hydrogen bonding pattern[26,40]. These findings suggest that the Hsp70 groove accommodates a degree of registry plasticity at the side-chain level while enforcing a conserved backbone geometry, providing a mechanistic basis for how diverse CMA-targeting motifs can be stably recognized without requiring identical residue alignments.

### Transient water molecules mediate interactions between CMA-targeting motifs and the SBDβ-lid interface of Hsc70

To delve deeper into the reason behind the stable binding of KFERQ-like peptides only in the presence of SBDα/lid, we evaluated the contribution of the surrounding water molecules to the binding stability. Our simulations suggested that water molecules mediate interactions between CMA-targeting motifs and the lid interface of Hsc70 via the formation of hydrogen bonds (Fig 5A, S7 Fig.). The time evolution of the number of water molecules bridging the substrate peptide and the lid domain indicates the involvement of ∼1-7 water molecules depending on the substrate. While the residues of substrate peptide and/or lid domain of Hsc70 involved in hydrogen bond formation is not consistent among different substrates and/or different time frames, interestingly, the water bridge was observed to be formed between the hydrophilic residues of lid domain with hydrophilic amino acid of KFERQ-like motifs (S8 Fig.).

**Fig 5.**
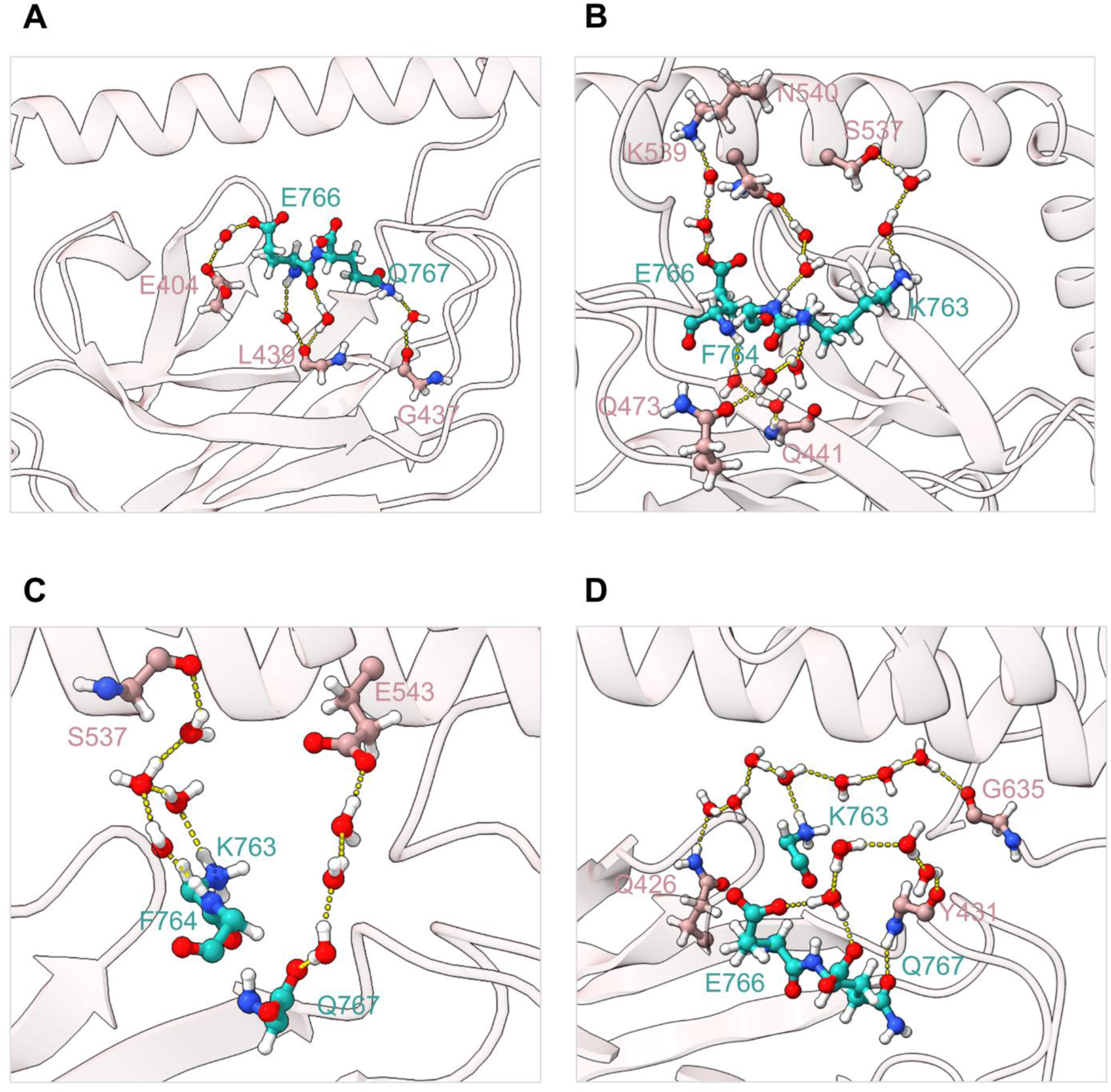
Water molecules mediate interactions between CMA-targeting motifs and the SBDβ-lid interface of Hsc70. Representative examples showcasing CMA-targeting motifs in Abeta interacting with the lid domain of Hsc70 through following number of water molecules: (A) one (B) two (C) three and (D) four. Hsc70 is represented in rosy brown and the interacting residues of CMA-targeting motifs in light sea green color.

In addition to the formation of a direct water bridge, extended water chains (two concurrent water bridges, or in some cases, three or more water molecules) where a substrate-water hydrogen bond is relayed through additional water molecules before reaching the lid domain, were also observed between the substrate peptide and the lid domain of Hsc70 (Fig 5). For instance, residue E766 of Abeta and residue K539 of SBDα were connected by two water molecules via hydrogen bonds (Fig 5B), whereas three water molecules mediate interactions between residue Q767 of Abeta and residue E543 of SBDα (Fig 5C). On similar lines, four water molecules bridges residue K763 of Abeta and residue G635 of SBDα (Fig 5D), further reinforcing the interaction network. Similar to the above observations, these bonds are not consistent in different substrates and/or in different time frames. We speculate that these transient hydrogen bonds dynamically stabilize the closed conformation of Hsc70.

Furthermore, we explored the contribution of water in stabilizing the interactions between the substrate peptide and SBDβ of Hsc70. The time evolution of the number of direct water bridges between the substrate peptide and the SBDβ indicates the involvement of ∼1-13 water molecules, (S9 Fig.). Overall, these results suggest that the role of water in CMA substrate recognition may be indirect but functionally significant, particularly in shaping the conformational landscape of the binding groove.

To summarize, the current work proposed two potential binding sites in Hsc70 for CMA substrates. The first site resides in the hydrophobic cavity inside the lid domain of Hsc70 encompassing residues in α1 and α3 helix. Notably, CMA-targeting motifs alone are insufficient to stabilize the interactions and that adjacent hydrophobic residues play a crucial role in reinforcing it. The second site was observed to be the canonical hydrophobic cavity inside the SBDβ domain of Hsc70. The binding in the site remained stable exclusively in the closed conformation of Hsc70, where transient water molecules mediated interactions between hydrophilic residues of the CMA-targeting motifs and the lid domain. Both identified sites have supporting interaction evidence in the literature[32,36]. However, whether our simulation results reflect a metastable intermediate or an alternative, physiologically relevant interaction pose remains to be validated. Further structural and biochemical analyses will be required to determine whether this modeled conformation corresponds to a functional state of CMA substrate recognition.

## Methods

### Preparation of Starting Inputs for Hsc70-Substrate Complex Generation

Fifteen experimentally verified CMA substrates were selected based on the following criteria: i) they are of human origin and ii) each contains a single canonical KFERQ-like motif. The presence and sequence of KFERQ-like motifs in these substrates were identified using the KFERQ finder V0.8 tool[13]. The sequences of these canonical motifs along with the full-length Hsc70 sequence and ADP, were used as an input to AlphaFold 3 (AF3)[29] to generate models of the binding complexes between Hsc70 and their respective CMA substrates. For truncated Hsc70 models comprising only the substrate-binding domain α (SBDα), residues 510-646 were used as input. Among the five models generated for each complex, the one with the highest interface predicted TM-score (iPTM) was selected for further analysis (S3 Table). The secondary structures of the substrate proteins were determined using the STRIDE[41] program, applied to structures obtained from the Protein Data Bank (PDB) and AlphaFold database[42–44]. To construct complexes of extended ‘KFERQ-like’ motifs with Hsc70, the following steps were performed. For each substrate protein, residues constituting the ‘KFERQ-type’ motif were first located in the sequence and mapped onto their corresponding positions in the 3D structure. The residues immediately flanking the motif on both N- and C-terminal sides were then analyzed for their secondary structure annotation. Adjacent residues were defined as variable-length stretches of amino acids contiguous to the KFERQ-type motif that (i) adopt the same secondary structure as the motif (in cases where the motif resides within an α-helix or β-sheet) or (ii) belong to an adjacent secondary structure element (in cases where the motif is located within a loop region). These residues, together with the KFERQ-type motif, were grouped and termed as ‘extended motifs’. The sequence of these extended motifs was fed as an input to AF3, along with truncated Hsc70 sequence (only SBDα domain, residues 510 to 646) to generate complex models. As before, the models with the highest iPTM scores were selected for further analysis.

Complexes of the KFERQ-like motifs from RND3, α-synuclein, and PARK7 with the closed conformation of full-length Hsc70 were constructed using a hybrid modeling approach. Initially, the interaction interfaces between each KFERQ-type motif and the AlphaFold 3 (AF3)-predicted open conformation of Hsc70 (which includes ADP though) were identified and used as reference contact maps. These contact interfaces were then preserved while fitting the respective motifs into the closed conformation of Hsc70, obtained from the AF3-predicted complex of Hsc70 with the KFERQ-like motif of Abeta. This procedure ensured that the substrate-Hsc70 interactions observed in the open state were retained within the closed-state framework of Hsc70 for subsequent analyses.

### Molecular Dynamics Simulations

To explore the stability of protein-protein interactions in the Hsc70-substrate complexes predicted using AlphaFold 3, all-atom MD simulations were performed using the GROMACS 2020.4 package for each model of Hsc70-CMA substrate complexes[45]. CHARMM36 force field was used to simulate protein complexes[46,47]. Preparation of initial systems was done by assigning standard protonation states to the ionizable residues corresponding to pH 7 using PropKa[48]. Cubic box with a gap of 10 Å between the surface of the protein complex and the edge was defined. Systems were solvated using TIP3P water model and neutralized by NaCl[49]. Energy minimization was done using Steepest Descent algorithm until the energy difference between steps was less than 1000 kJ mol^-1^ nm^-1^[50]. Two-step equilibration was performed, first using the NVT ensemble, followed by the NPT ensemble. The temperature was controlled to 300 K by a V-rescale thermostat[51] and pressure to 1 bar using the Parrinello–Rahman barostat[52]. A time step of 2 fs was used and at every 100 steps coordinates were written. Amber as well as OPLS force fields were used for two complexes (α-Synuclein and PARK7) to analyze the effect of force fields[53,54]. The analysis was carried out either using in-build gromacs utilities or in-house scripts.

## Acknowledgements

N.M. acknowledges the financial support from the Shiv Nadar Foundation and the DST Inspire Faculty Grant [DST/INSPIRE/04/2022/001599]. D.S. acknowledges the junior research fellowship from the Shiv Nadar Foundation. We are grateful to the supercomputing facility, Magus02, at SNIoE for the compute support.

## Supplementary information Captions

**S1 Table. List of experimentally verified CMA substrates in humans along with the canonical CMA-targeting motif in them.** The superscript numbers indicate the starting position of the CMA-targeting motif.

**S1 Fig. CMA-targeting motifs in substrate proteins stably binds to the lid/SBDα of Hsc70 only in the presence of extended/adjacent flanking residues.**

(A) Time evolution of the minimum distance between Hsc70 SBDα and the KFERQ-like motif of 3 different substrates. (B) Time evolution of the minimum distance between HSC70 SBDα and the extended KFERQ-like motif of 15 different CMA substrates.

**S2 Fig. Extended CMA-targeting motifs in substrate proteins display promiscuous orientation in the binding hydrophobic cavity of Hsc70, with major energetic contributions from hydrophobic residues in substrate protein.**

(A) Orientation of CMA substrate peptides (CMA-targeting motifs along with flanking residues or extended CMA-targeting motifs) within the hydrophobic cavity of Hsc70 at 1μs of MD simulations, with the substrate segment depicted in gray cartoon representation. (B) Residue-wise decomposition of binding energies of extended CMA substrates peptides. Only values less than - 2.5 kcal/mol are shown for clarity.

**S3 Fig. Interacting residue pairs between Hsc70 and extended CMA-targeting motifs differ across substrates, suggesting a flexible and promiscuous mode of binding.** The plots depict the minimum distance between representative residue pairs with binding energies less than −2.5 kcal/mol.

**S4 Fig. Promiscous mode of binding of extended CMA-targeting motifs in the hydrophobic cavity of Hsc70’s lid domain was observed in presence of different force fields**

(A) Snapshot at 1μs of MD simulations depicting orientation of extended CMA-targeting motifs in two substrate peptides within the hydrophobic cavity of Hsc70 using three different force fields (Amber: rosy brown, OPLS:dark khaki, and CHARMM36:cornflower blue), residues with binding energies less than −2.5 kcal/mol are highlighted in the ball and stick model. Residues which are part of CMA-targeting motif(s) are labelleled in red color while the flanking residues are labelled in black. (B) Residue-wise decomposition of binding energies of PARK7 and α-Synuclein using three different force fields (Amber, OPLS, and CHARMM36), with different values in each force field indicating the flexibility of substrate binding modes.

**S5 Fig. AlphaFold 3 predicted complexes of Hsc70 and CMA-targeting motifs of fifteen shortlisted CMA substrates suggests SBDβ to be the possible binding site.**KFERQ-like motifs (light sea green) are bound to hydrophobic cleft of SBDβ (dark gray) in two hsc70 conformations i.e. open and closed(*). For Clarity, only the SBDβ domain is shown.

**S2 Table. Segregation of residues within CMA-targeting motifs into distinct binding pockets (labelling as per previous literature**[37,38]**) based on their stable interaction sites.** Data are shown for seven closed Hsc70–substrate complexes that remained stable during the MD simulations.

**S6 Fig. Backbone of CMA-targeting motifs in substrate proteins are involved in interaction with backbone of residues in SBDβ of Hsc70, suggesting preservation of backbone alignment in the interaction mode of the substrate.** Figure depicts hydrogen bonding (yellow color) between backbone of CMA-targeting motif in seven CMA substrates (Abeta, AnnexinA1, PEA-15, HBB, RND3, α-Synuclein and PARK7) with backbone of two key residues (having >90% occupancy) in SBDβ viz. Ala406 and Thr429. The interacting residues are explicitly labelled.

**S7 Fig. Water-mediated hydrogen bonds stabilize the binding between the lid domain of Hsc70 and the CMA-targeting motif in the substrate protein.**

**(A)** The time evolution of the number of water molecules forming hydrogen bonds between the CMA-targeting motif in different substrate proteins and the lid domain of Hsc70 for four stable complexes in the MD simulations with Hsc70 in the closed conformation. (**B)** The time evolution of the number of water molecules forming hydrogen bonds between the CMA-targeting motif in different substrate proteins and the lid domain of Hsc70 for three stable closed complexes in the MD simulations with Hsc70 which were initially predicted in the open conformation by AF3.

**S8 Fig. Transient water molecules stabilizes the binding of the substrate peptides with the lid domain of Hsc70 at different time frames.** Snapshots of seven closed stable complexes forming water mediated hydrogen bonding between lid domain and CMA-targeting motifs at three different time frames. Interacting residues of Hsc70 lid domain (rosy brown) and CMA-targeting motifs (light sea green), along with water molecules are represented in ball-stick form.

**S9 Fig. Water-mediated hydrogen bonds stabilize the binding between the SBDβ domain of Hsc70 and the CMA-targeting motif in the substrate protein.**

(A) Time evolution of the number of water molecules forming hydrogen bonds between the CMA-targeting motif in different substrate proteins and SBDβ domain of Hsc70 for four stable complexes in the MD simulations, with Hsc70 in the closed conformation. (B) Time evolution of the number of water molecules forming hydrogen bonds between the CMA-targeting motif in different substrate proteins and the SBDβ domain of Hsc70 for three stable complexes in the MD simulations with Hsc70 predicted in the open conformation and converted to closed conformation.

**S3 Table. List of complexes generated using AlphaFold 3 utilizing segments of Hsc70 and substrate proteins, along with their confidence scores (iptm and ptm).**

## Data availability statement

The data files related to this study have been made available at: https://github.com/CSBLabNM/Project_HSC70.git

