## Supplementary Information for "Investigation of CMA Substrate Interactions with Hsc70 Reveals Potential Auxiliary Binding Sites"

Devid Sahu[1], Nidhi Malhotra[1]*

[1] Department of Chemistry, School of Natural Sciences, Shiv Nadar Institution of Eminence, Gautam Buddha Nagar, Uttar Pradesh, India

*Corresponding author

**S1 Table.** List of experimentally verified CMA substrates in humans along with the canonical CMA-targeting motif in them. The superscript numbers indicate the starting position of the CMA-targeting motif.

| CMA Substrates | CMA-targeting motif |
|---|---|
| Rho-related GTP-binding protein RhoE | $^{69}$QRIEL |
| Amyloid-beta precursor protein | $^{763}$KFFEQ |
| Annexin A1 | $^{6}$EFLKQ |
| Annexin A2 | $^{226}$QKVFD |
| Annexin A6 | $^{564}$QEFIK |
| Lysosomal acid glucosylceramidase | $^{295}$QRDFI |
| G1/S-specific cyclin-D1 | $^{49}$QKEVL |
| Astrocytic phosphoprotein PEA-15 | $^{110}$DIIRQ |
| Hemoglobin subunit beta | $^{40}$QRFFE |
| Serine/threonine-protein kinase Chk1 | $^{336}$DKLVQ |
| α-Synuclein | $^{95}$VKKDQ |
| Annexin A4 | $^{93}$QELRR |
| Major prion protein | $^{208}$RVVEQ |
| Pyruvate kinase PKM | $^{227}$QDLKF |
| Parkinson disease protein 7 | $^{91}$ILKEQ |

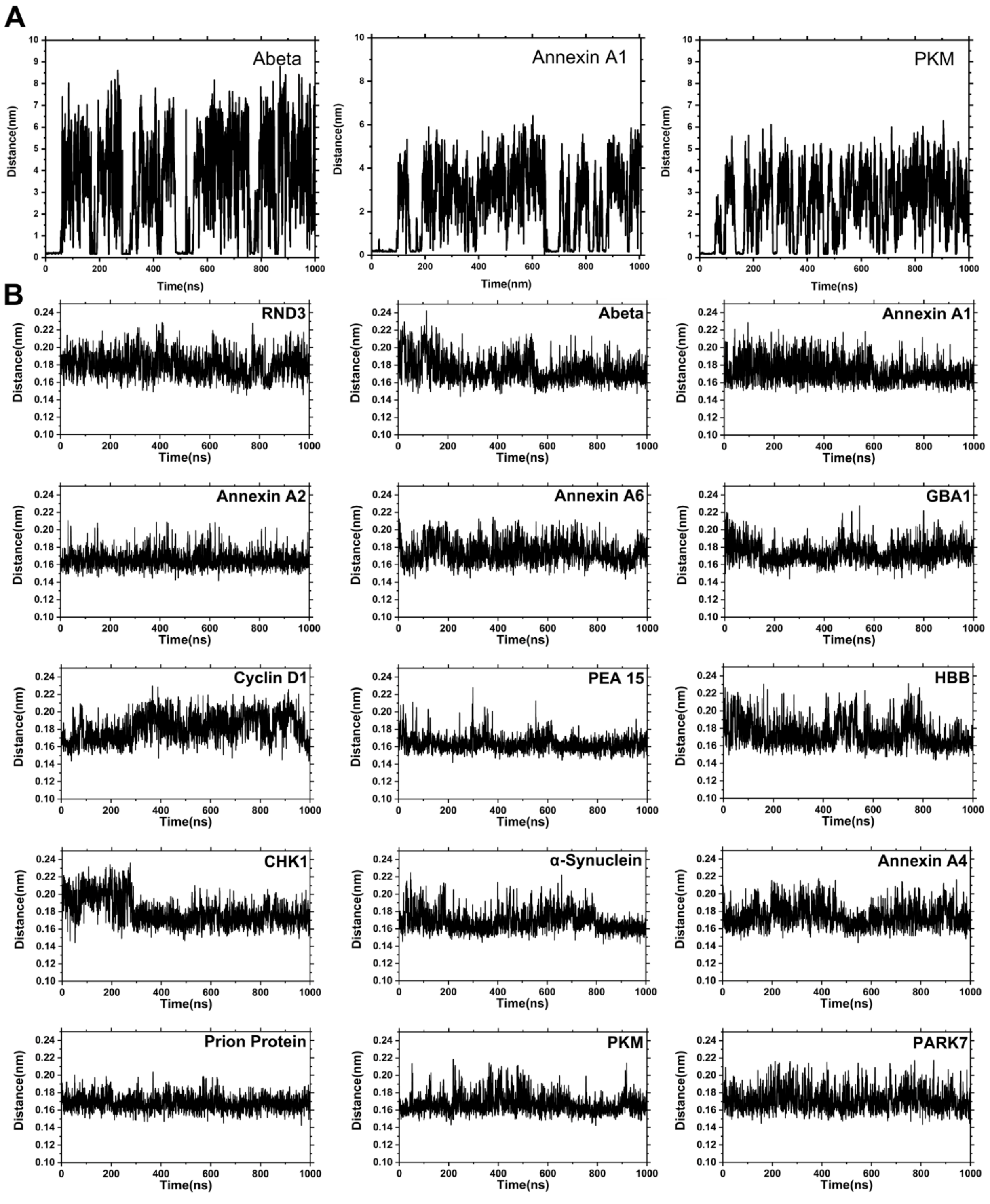


**S1 Fig. CMA-targeting motifs in substrate proteins stably binds to the lid/SBDα of Hsc70 only in the presence of extended/adjacent flanking residues.**

(A) Time evolution of the minimum distance between Hsc70 SBDα and the KFERQ-like motif of 3 different substrates. (B) Time evolution of the minimum distance between HSC70 SBDα and the extended KFERQ-like motif of 15 different CMA substrates.

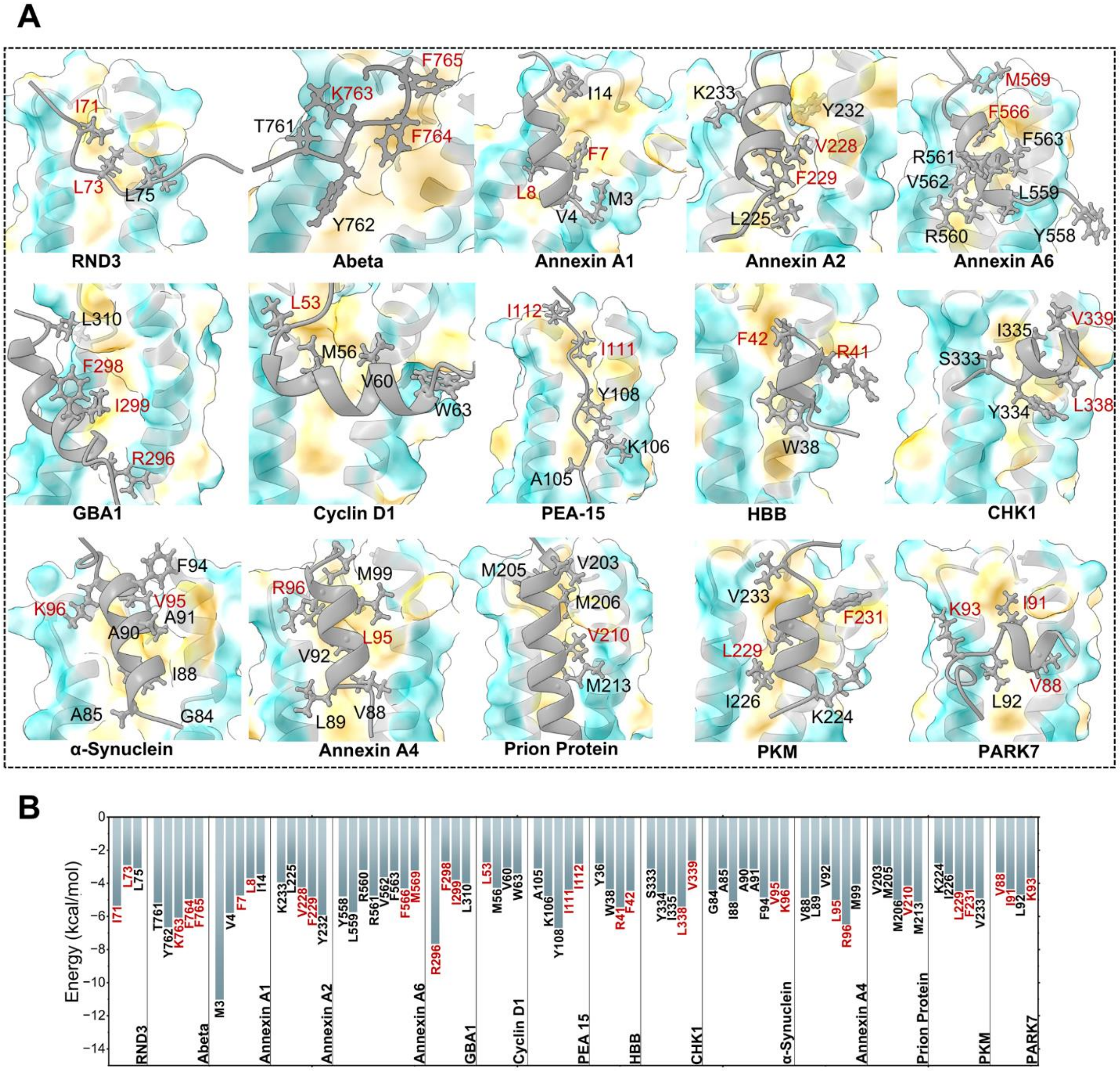


**S2 Fig. Extended CMA-targeting motifs in substrate proteins display promiscuous orientation in the binding hydrophobic cavity of Hsc70, with major energetic contributions from hydrophobic residues in substrate protein.**

(A) Orientation of CMA substrate peptides (CMA-targeting motifs along with flanking residues or extended CMA-targeting motifs) within the hydrophobic cavity of Hsc70 at 1μs of MD simulations, with the substrate segment depicted in gray cartoon representation. (B) Residue-wise decomposition of binding energies of extended CMA substrates peptides. Only values less than -2.5 kcal/mol are shown for clarity.

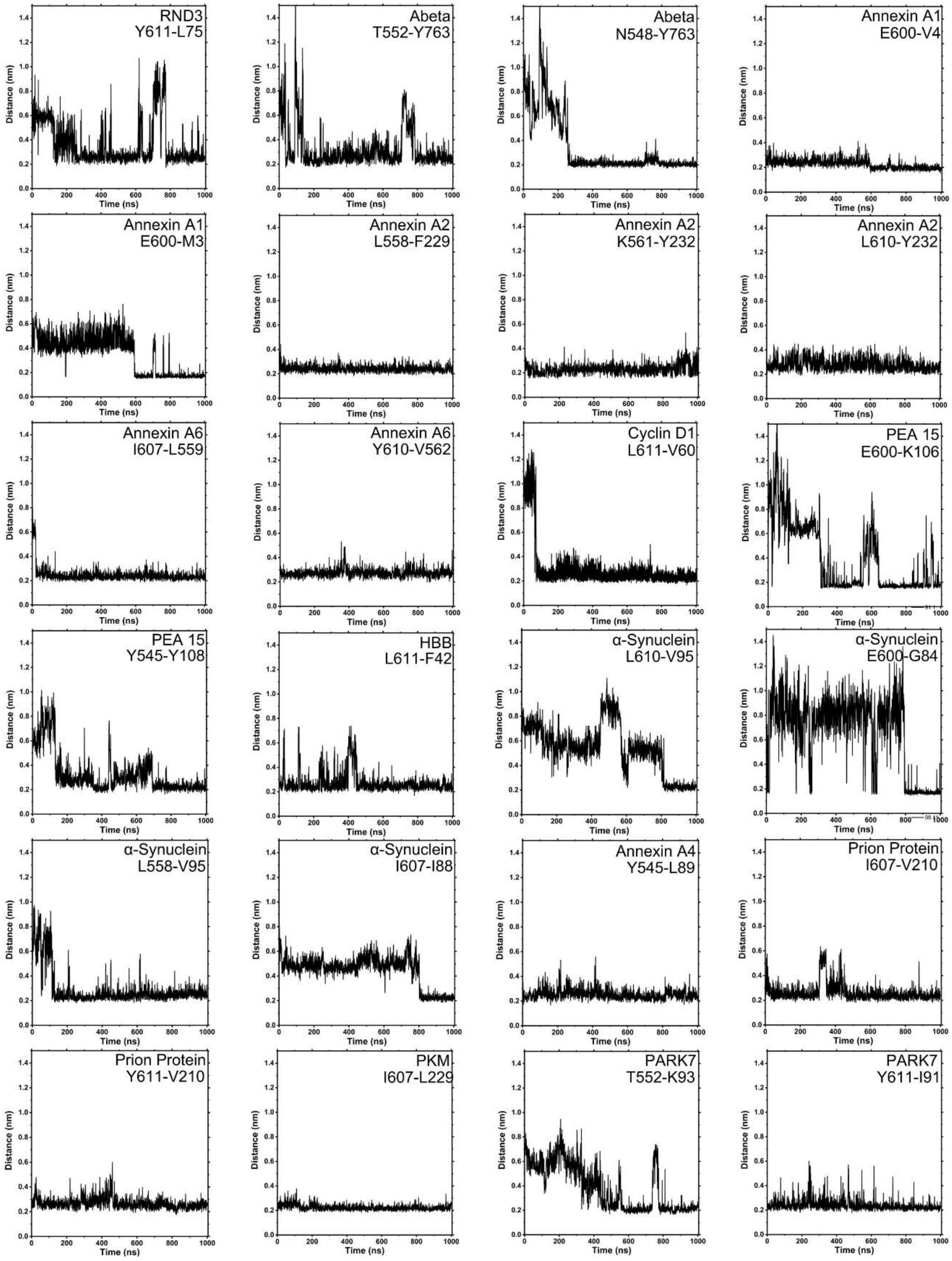


**S3 Fig. Interacting residue pairs between Hsc70 and extended CMA-targeting motifs differ across substrates, suggesting a flexible and promiscuous mode of binding.** The plots depict the minimum distance between representative residue pairs with binding energies less than -2.5 kcal/mol.

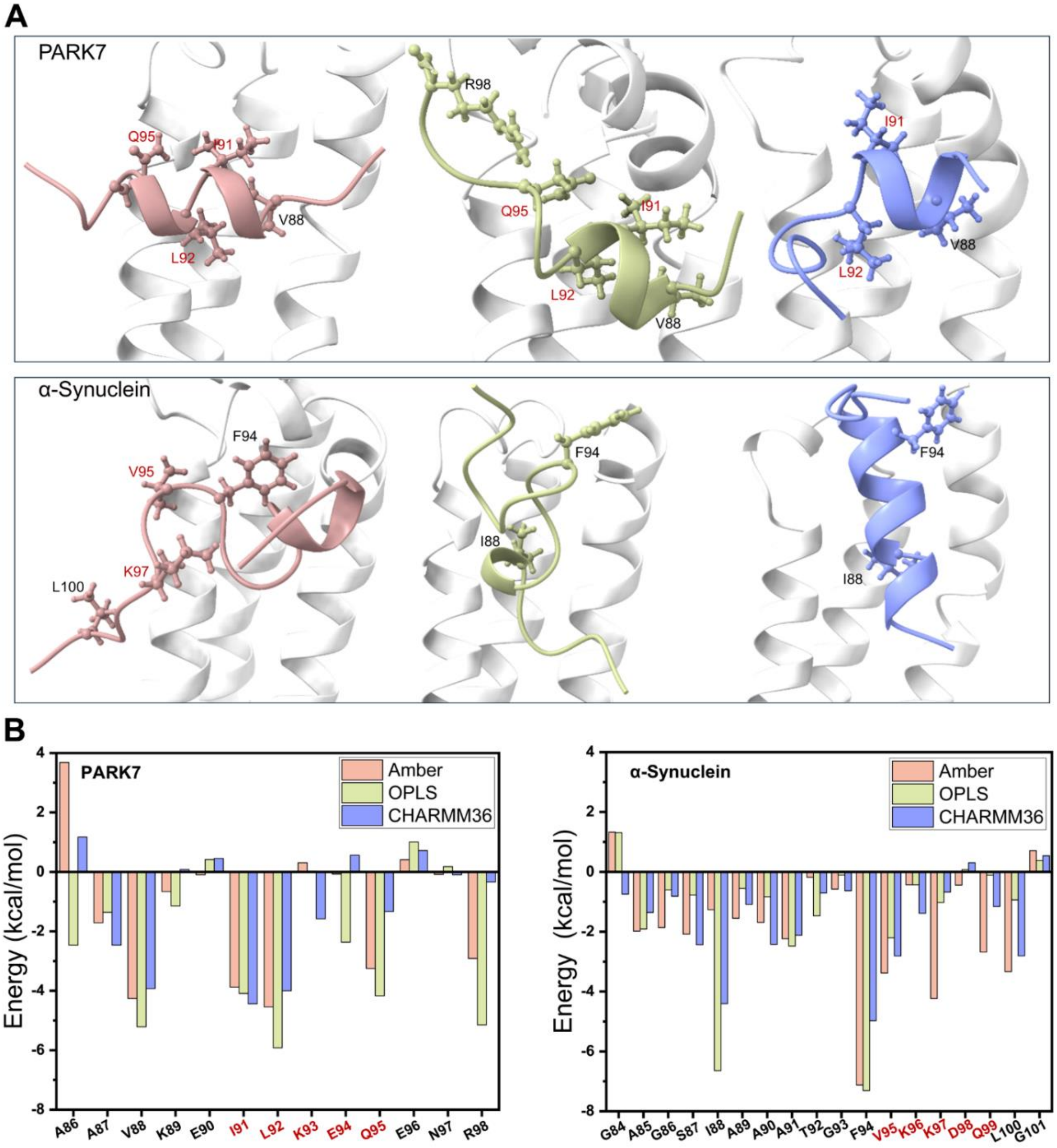


**S4 Fig. Promiscous mode of binding of extended CMA-targeting motifs in the hydrophobic cavity of Hsc70's lid domain was observed in presence of different force fields.**
(A) Snapshot at 1μs of MD simulations depicting orientation of extended CMA-targeting motifs in two substrate peptides within the hydrophobic cavity of Hsc70 using three different force fields (Amber: rosy brown, OPLS:dark khaki, and CHARMM36:cornflower blue), residues with binding energies less than -2.5 kcal/mol are highlighted in the ball and stick model. Residues which are part of CMA-targeting motif(s) are labelleled in red color while the flanking residues are labelled in black. (B) Residue-wise decomposition of binding energies of PARK7 and α-Synuclein using three different force fields (Amber, OPLS, and CHARMM36), with different values in each force field indicating the flexibility of substrate binding modes.

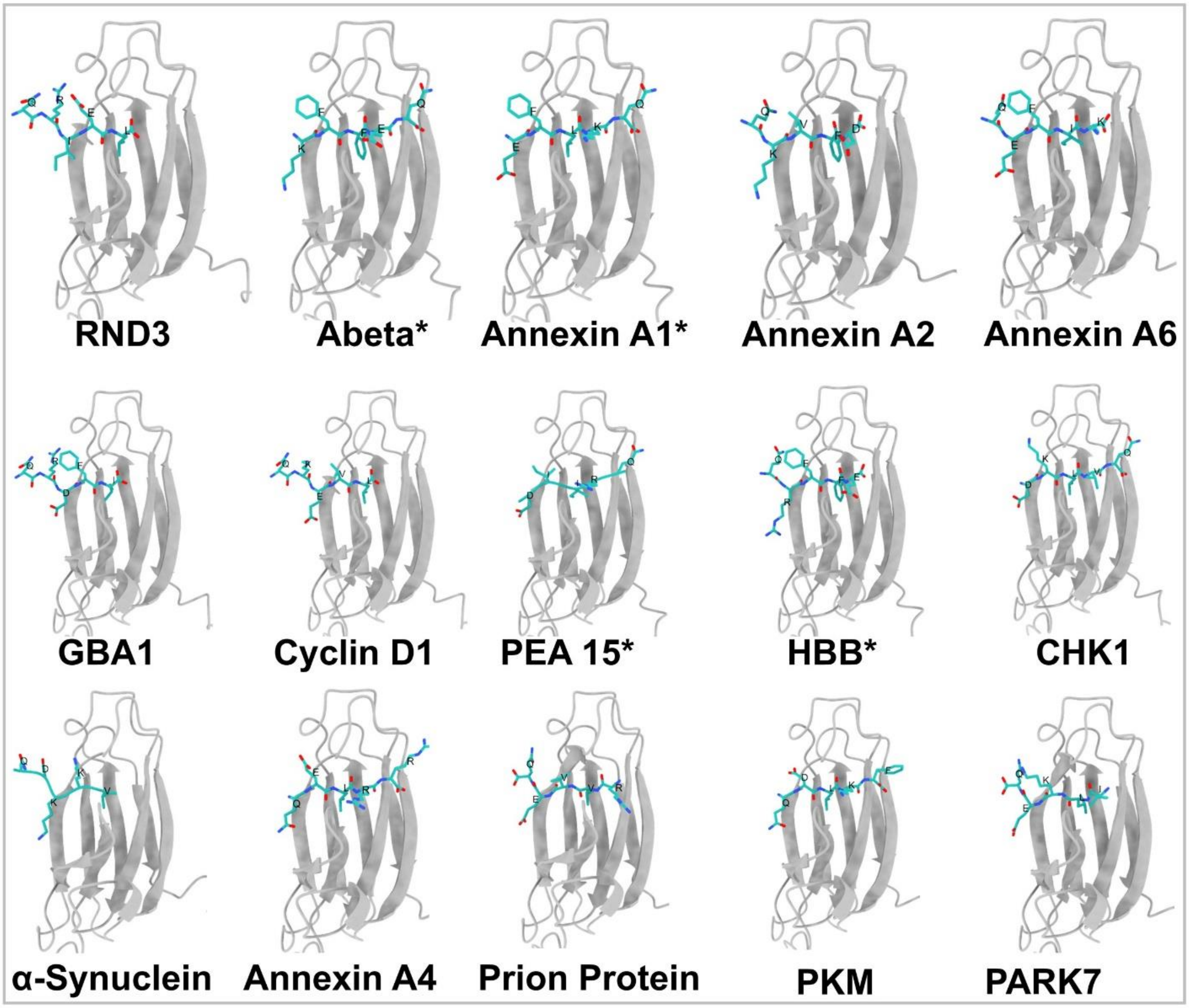


**S5 Fig. AlphaFold 3 predicted complexes of Hsc70 and CMA-targeting motifs of fifteen shortlisted CMA substrates suggests SBDβ to be the possible binding site.**
KFERQ-like motifs (light sea green) are bound to hydrophobic cleft of SBDβ (dark gray) in two hsc70 conformations i.e. open and closed(*). For Clarity, only the SBDβ domain is shown.

**S2 Table.** Segregation of residues within CMA-targeting motifs into distinct binding pockets (labelling as per previous literature[1,2]) based on their stable interaction sites. Data are shown for seven closed Hsc70–substrate complexes that remained stable during the MD simulations.

| **Substrate** | **-4** | **-3** | **-2** | **-1** | **0** | **+1** | **+2** | **+3** |
|---|---|---|---|---|---|---|---|---|
| Abeta | | | K | F | F | E | Q | |
| Annexin A1 | | | E | F | L | K | Q | |
| PEA 15 | | | D | I | I | R | Q | |
| HBB | | Q | R | F | F | E | | |
| RND3 | Q | R | I | E | L | | | |
| α-Synuclein | Q | D | K | K | V | | | |
| PARK7 | | Q | E | K | L | I | | |

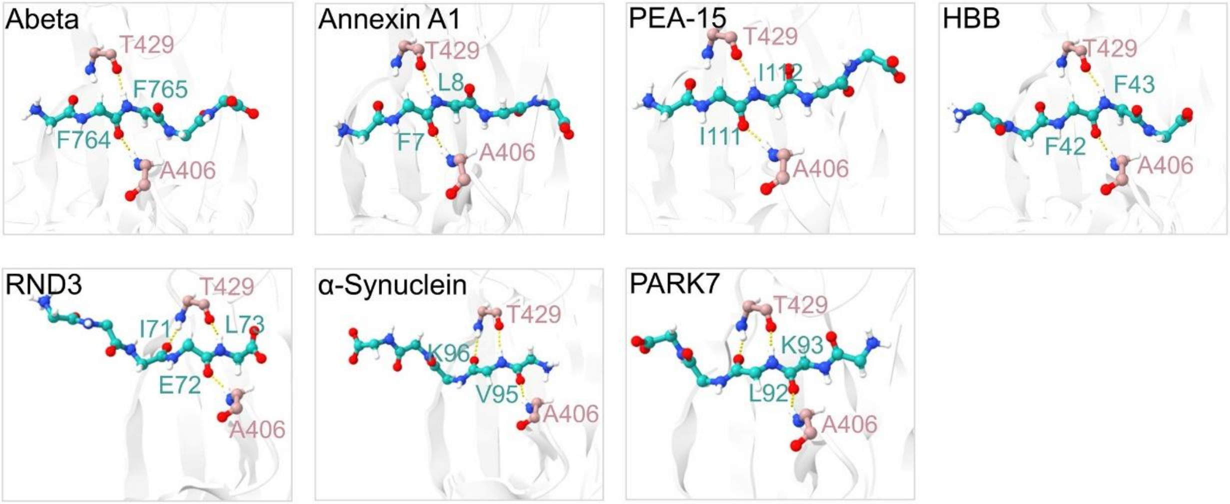


**S6 Fig. Backbone of CMA-targeting motifs in substrate proteins are involved in interaction with backbone of residues in SBDβ of Hsc70, suggesting preservation of backbone alignment in the interaction mode of the substrate.**

Figure depicts hydrogen bonding (yellow color) between backbone of CMA-targeting motif in seven CMA substrates (Abeta, AnnexinA1, PEA-15, HBB, RND3, α-Synuclein and PARK7) with backbone of two key residues (having >90% occupancy) in SBDβ viz. Ala406 and Thr429. The interacting residues are explicitly labelled.

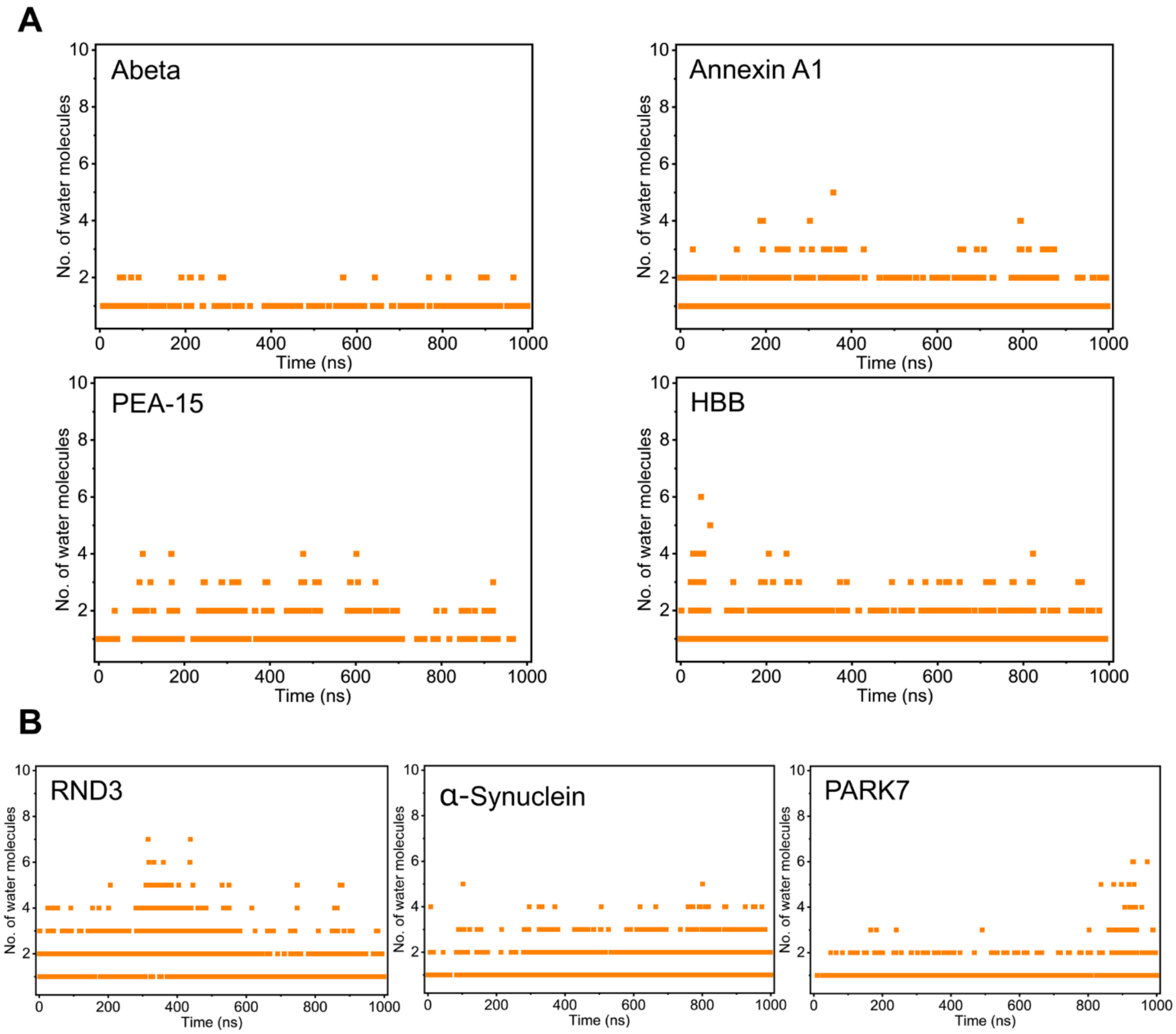


**S7 Fig. Water-mediated hydrogen bonds stabilize the binding between the lid domain of Hsc70 and the CMA-targeting motif in the substrate protein.**

(A) The time evolution of the number of water molecules forming hydrogen bonds between the CMA-targeting motif in different substrate proteins and the lid domain of Hsc70 for four stable complexes in the MD simulations with Hsc70 in the closed conformation. (B) The time evolution of the number of water molecules forming hydrogen bonds between the CMA-targeting motif in different substrate proteins and the lid domain of Hsc70 for three stable closed complexes in the MD simulations with Hsc70 which were initially predicted in the open conformation by AF3.

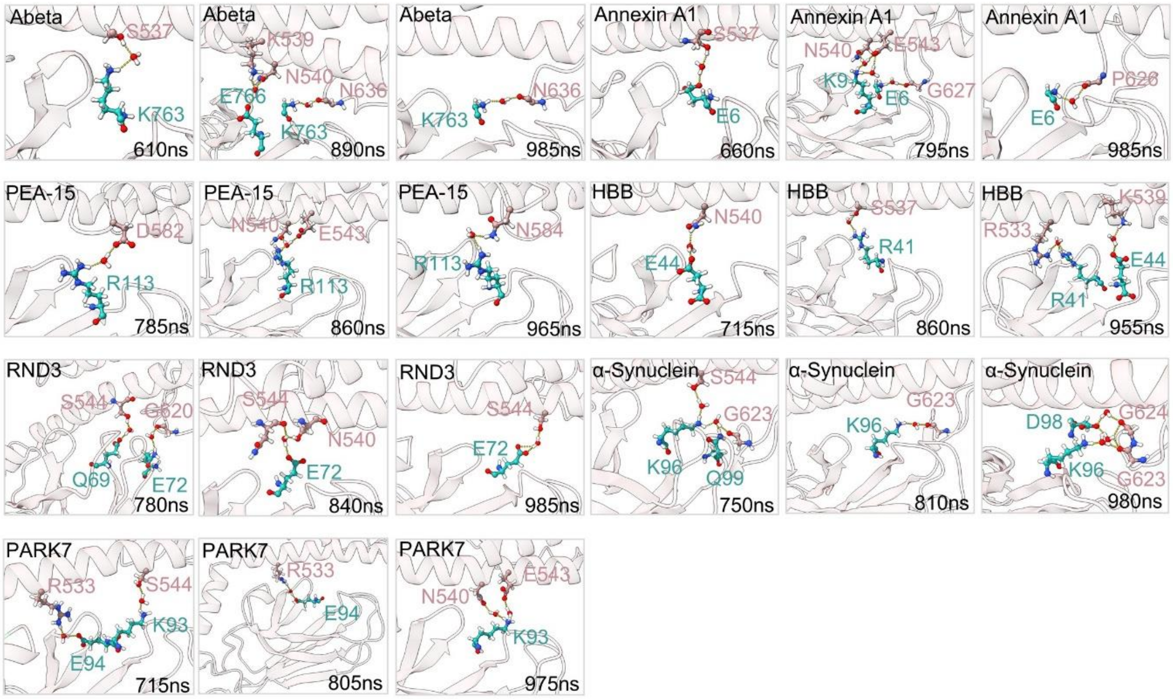


**S8 Fig. Transient water molecules stabilizes the binding of the substrate peptides with the lid domain of Hsc70 at different time frames.**

Snapshots of seven closed stable complexes forming water mediated hydrogen bonding between lid domain and CMA-targeting motifs at three different time frames. Interacting residues of Hsc70 lid domain (rosy brown) and CMA-targeting motifs (light sea green), along with water molecules are represented in ball-stick form.

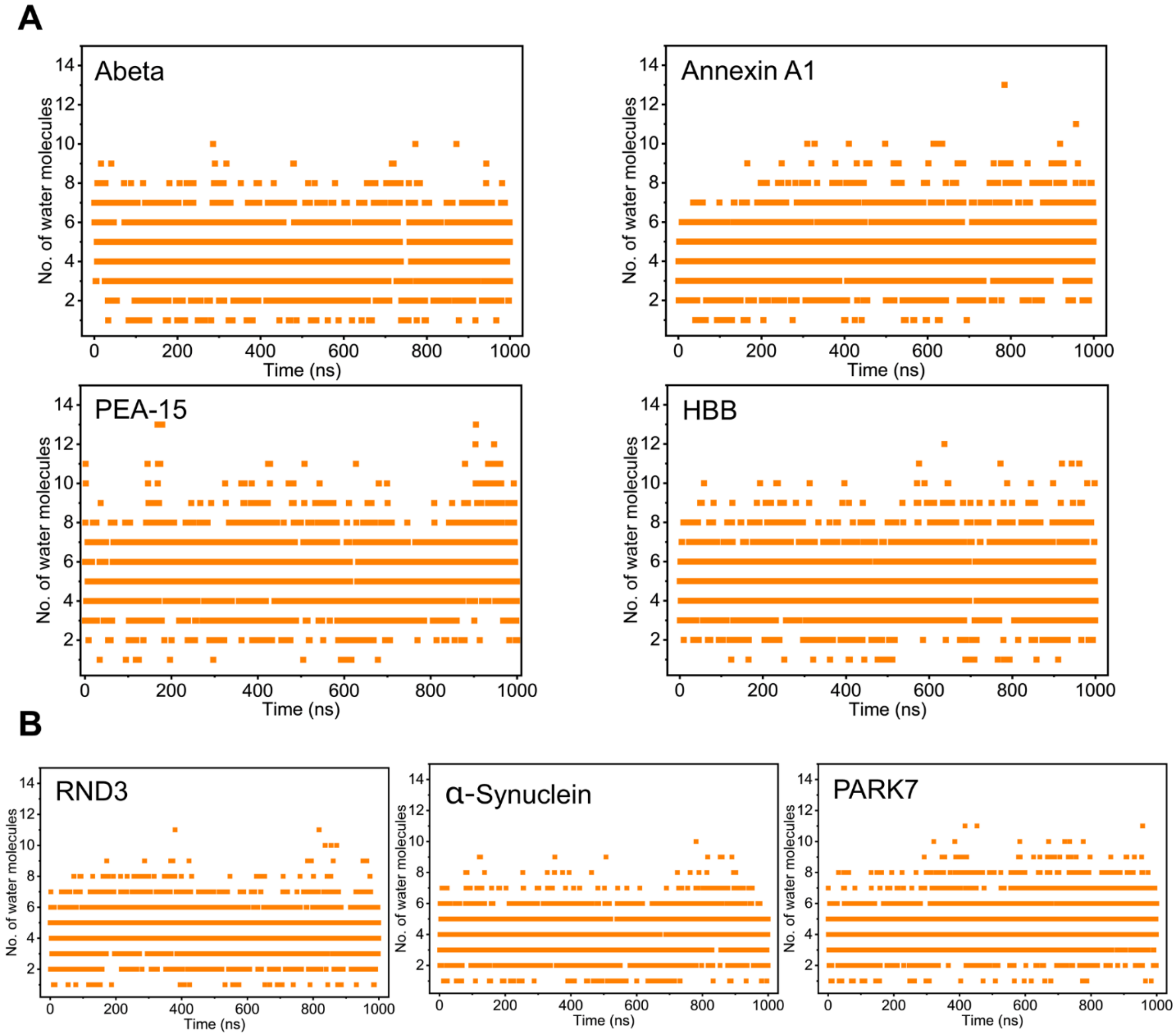


**S9 Fig. Water-mediated hydrogen bonds stabilize the binding between the SBDβ domain of Hsc70 and the CMA-targeting motif in the substrate protein.**

(A) Time evolution of the number of water molecules forming hydrogen bonds between the CMA-targeting motif in different substrate proteins and SBDβ domain of Hsc70 for four stable complexes in the MD simulations, with Hsc70 in the closed conformation. (B) Time evolution of the number of water molecules forming hydrogen bonds between the CMA-targeting motif in different substrate proteins and the SBDβ domain of Hsc70 for three stable complexes in the MD simulations with Hsc70 predicted in the open conformation and converted to closed conformation.

**S3 Table.** List of complexes generated using AlphaFold 3 utilizing segments of Hsc70 and substrate proteins, along with their confidence scores (iptm and ptm).

| Part of Hsc70 | CMA Substrates | Substrates Name | iptm | ptm |
|---|---|---|---|---|
| Lid domain | KFERQ-like motif | Rho-related GTP-binding protein RhoE | 0.36 | 0.66 |
| | | Amyloid-beta precursor protein | 0.39 | 0.66 |
| | | Annexin A1 | 0.28 | 0.64 |
| | | Annexin A2 | 0.4 | 0.66 |
| | | Annexin A6 | 0.36 | 0.66 |
| | | Lysosomal acid glucosylceramidase | 0.33 | 0.66 |
| | | G1/S-specific cyclin-D1 | 0.29 | 0.65 |
| | | Astrocytic phosphoprotein PEA-15 | 0.41 | 0.66 |
| | | Hemoglobin subunit beta | 0.4 | 0.65 |
| | | Serine/threonine-protein kinase Chk1 | 0.34 | 0.65 |
| | | α-synuclein | 0.29 | 0.65 |
| | | Annexin A4 | 0.32 | 0.66 |
| | | Major prion protein | 0.33 | 0.66 |
| | | Pyruvate kinase PKM | 0.32 | 0.66 |
| | | Parkinson disease protein 7 | 0.24 | 0.65 |
| | KFERQ-like motif with extended amino acids | Rho-related GTP-binding protein RhoE | 0.38 | 0.63 |
| | | Amyloid-beta precursor protein | 0.4 | 0.64 |
| | | Annexin A1 | 0.44 | 0.63 |
| | | Annexin A2 | 0.48 | 0.64 |
| | | Annexin A6 | 0.42 | 0.63 |
| | | Lysosomal acid glucosylceramidase | 0.53 | 0.64 |
| | | G1/S-specific cyclin-D1 | 0.27 | 0.58 |
| | | Astrocytic phosphoprotein PEA-15 | 0.57 | 0.67 |
| | | Hemoglobin subunit beta | 0.42 | 0.64 |
| | | Serine/threonine-protein kinase Chk1 | 0.51 | 0.65 |
| | | α-synuclein | 0.53 | 0.64 |
| | | Annexin A4 | 0.47 | 0.63 |
| | | Major prion protein | 0.3 | 0.58 |
| | | Pyruvate kinase PKM | 0.57 | 0.65 |
| | | Parkinson disease protein 7 | 0.51 | 0.65 |
| Full length Hsc70 with ADP | KFERQ-like motif | Rho-related GTP-binding protein RhoE | 0.85 | 0.66 |
| | | Amyloid-beta precursor protein | 0.84 | 0.62 |
| | | Annexin A1 | 0.85 | 0.63 |
| | | Annexin A2 | 0.85 | 0.65 |
| | | Annexin A6 | 0.85 | 0.64 |
| | | Lysosomal acid glucosylceramidase | 0.85 | 0.66 |
| | | G1/S-specific cyclin-D1 | 0.85 | 0.66 |
| | | Astrocytic phosphoprotein PEA-15 | 0.85 | 0.63 |
| | | Hemoglobin subunit beta | 0.85 | 0.63 |
| | | Serine/threonine-protein kinase Chk1 | 0.85 | 0.66 |
| | | α-synuclein | 0.89 | 0.73 |
| | | Annexin A4 | 0.85 | 0.67 |
| | | Major prion protein | 0.85 | 0.66 |
| | | Pyruvate kinase PKM | 0.86 | 0.66 |
| | | Parkinson disease protein 7 | 0.85 | 0.66 |
| Full length Hsp70 with ADP | NRLLLTG peptide | NRLLLTG peptide | 0.81 | 0.63 |
